# Isolation of a putative sulfur comproportionating microorganism

**DOI:** 10.1101/2023.06.08.544259

**Authors:** Heidi S. Aronson, Douglas E. LaRowe, Jennifer L. Macalady, Jan P. Amend

## Abstract

Sulfur comproportionation is a heretofore undiscovered microbial catabolism that was predicted based on thermodynamic calculations. Here, we report the isolation of an *Acidithiobacillus thiooxidans* strain from extremely low pH snottite biofilms in the karst at Frasassi, Italy. The strain grew to cell densities of >10^7^ cells mL^-1^ in autotrophic sulfur comproportionation medium. Whole genome sequencing of the isolate revealed the presence of numerous genes involved in sulfur transformations that could be linked in a sulfur comproportionation pathway. We describe an experimental framework, including measurements of sulfate, sulfide, and S^0^ concentrations, electron microscopy, and stable and radioisotope incubations coupled with NanoSIMS, scintillation counting and isotope ratio mass spectrometry, for future searches of sulfur comproportionators.

**Significance:** The prediction of and search for novel microbial catabolic reactions can be streamlined by using thermodynamics to identify energy-yielding redox reactions that may be catalyzed by microorganisms. This strategy has been used to successfully predict several previously overlooked microbial catabolic reactions, including anaerobic ammonia oxidation (anammox), anaerobic oxidation of methane (AOM), and complete ammonia oxidation (comammox). Sulfur comproportionation, or the coupled reduction of sulfate and oxidation of sulfide to form elemental sulfur, was predicted by thermodynamic calculations to exist as a microbial catabolism in low pH, low-temperature environments. In this study, we describe the isolation of the first putative sulfur comproportionating microorganism and provide a detailed experimental approach that can be applied to future investigations of this novel link in the biogeochemical sulfur cycle.

## Main text

Sulfur comproportionation (Rxn 1), or the reaction of sulfide and sulfate to form elemental sulfur

(S^0^),

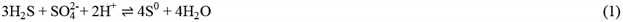

was recently proposed as a novel microbial catabolic reaction on the basis of Gibbs energy calculations (1). This reaction is exergonic in low-temperature, acidic environments with high sulfide and sulfate concentrations. Snottite biofilms within the Frasassi cave system (Italy) provide ideal conditions for sulfur comproportionation, with an *in situ* Gibbs free energy yield of 52 kJ mol^-1^.

Snottites were collected from Frasassi as inoculum for a custom medium targeted for the enrichment of autotrophic sulfur comproportionators. Incubation conditions were selected that yield the same amount of energy for sulfur comproportionation as the sample site. The enrichment was purified by dilution to extinction. The resulting isolate was identified as *Acidithiobacillus thiooxidans* with 99.86% sequence similarity to the type strain in the full-length 16S rRNA gene and was designated strain RS19-104. *A. thiooxidans*, an autotrophic, sulfur-oxidizing acidophilic bacterium, is abundant in Frasassi snottite biofilms, making up over 70% of the microbial community there (2), and is thought to be the dominant primary producer of the snottites. Cells of strain RS19-104 were rod-shaped and 1-2 μm in length (Fig. 1).

**Figure 1.**
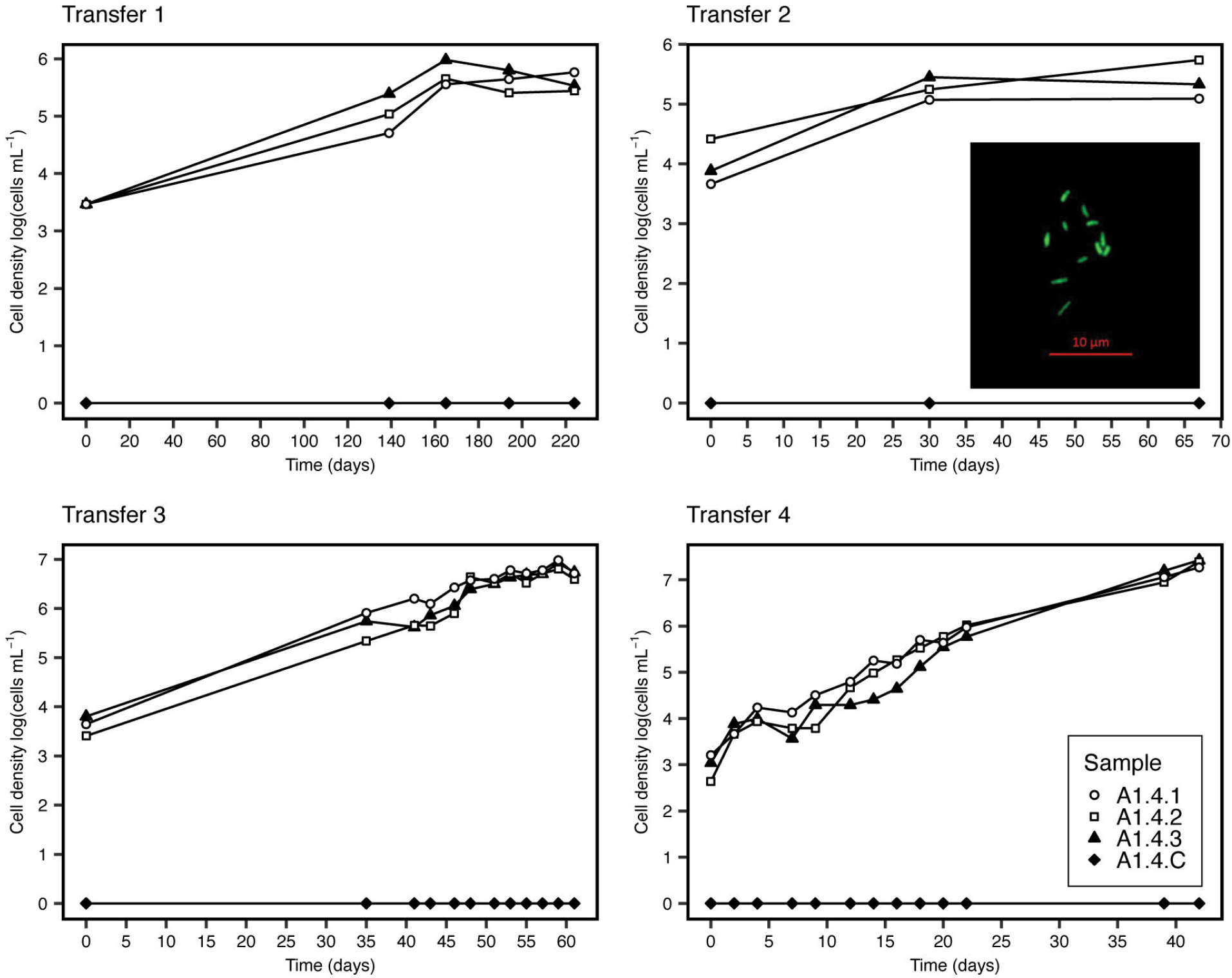
Cell densities in log (cells ml^-1^) of comproportionation cultures as a function of time. Each panel represents a separate transfer of three inoculated bottles (samples A1.X.1 – A1.X.3) with an uninoculated negative control bottle (1.X.C, black diamonds). The inset image in the “Transfer 2” panel shows cells of strain RS19-104 visualized under fluorescence microscopy. Scale bar represents 10 μm.

Cultures of strain RS19-104 (*n* = 12) showed increases in cell density of 2-3 orders of magnitude over the course of 1-2 months, with the highest cell densities reaching >10^7^ cells mL^-1^ (Fig. 1). Because *A. thiooxidans* can grow via aerobic oxidation of reduced sulfur compounds, possible oxygen contamination of the growth medium was considered. However, several lines of evidence suggest that strain RS19-104 was not growing as an aerobic sulfide oxidizer. Anaerobic technique was strictly applied with RS19-104 cultures, which included using blue butyl stoppers, adding a slight N_2_/CO_2_ overpressure in the headspace to prevent gas from leaking into the serum bottles, flushing needles with N_2_, working in an anaerobic chamber whenever possible, using resazurin as a redox indicator, and adding 2 mM sulfide as a reductant and electron donor. We therefore conclude that oxygen leakage cannot explain the growth of *A. thiooxidans* RS19-104 that we observed.

The draft genome of strain RS19-104 was 100% complete with 2.07% contamination and an average nucleotide identity and amino acid identity with *A. thiooxidans* ATCC 19377^T^ of 98.62% and 98.51%, respectively. Numerous genes encoding enzymes related to sulfur cycling were present in the genome. A sulfur comproportionating microorganism might use known sulfide oxidation and sulfate reduction pathways linked in previously unexpected ways. Because *A. thiooxidans* grows via aerobic sulfide oxidation using sulfide-quinone reductase (*sqr*), we postulated that Sqr might be used under comproportionation conditions to oxidize sulfide and donate electrons to the quinone pool. However, it was unknown whether strain RS19-104 could use sulfate as an electron acceptor. Several species of *Acidithiobacillus* are facultatively capable of anaerobic respiration using Fe(III) or S^0^ as electron acceptors (4–10), suggesting that anaerobic growth using sulfate as an electron acceptor is plausible. The presence of a complete dissimilatory sulfate reduction pathway (*sat, aprAB, dsrAB*) would indicate that strain RS19-104 uses sulfate for respiration. While *sat* was present in the genome, *aprAB* and *dsrAB* were absent. Instead, genes associated with the assimilatory sulfate reduction pathway, including *sat, cysC*, and *cysH* were present, suggesting that strain RS19-104 has the potential to produce sulfite from sulfate. The assimilatory sulfate reduction pathway consumes 2 ATP to convert sulfate into sulfide destined to make the S-bearing amino acids cysteine and methionine. However, the genome of strain RS19-104 also contains the gene *cysM*, which encodes an enzyme used to assimilate sulfide directly from the environment. Because strain RS19-104 grows in a sulfide-rich environment, it is plausible that it would use sulfide for biosynthesis rather than invest energy into sulfate assimilation. We propose that strain RS19-104 could be using the assimilatory sulfate reduction pathway for comproportionation. Alternatively, a gene encoding SoeABC, which oxidizes sulfite to sulfate while donating electrons to the quinone pool, was also found in the genome (11, 12). If acting in reverse, SoeABC could catalyze the reduction of sulfate to form sulfite.

Once sulfite is produced by sulfate reduction, there are several enzymes that, if acting in reverse, could catalyze the conversion of sulfite to S^0^. SoeABC was shown to operate in the reverse direction by reducing S^0^, thiosulfate, tetrathionate, and sulfite *in vitro* (12). While the bidirectionality of SoeABC was not confirmed *in vivo*, it is possible that it could reduce sulfite to S^0^ in strain RS19-104. A gene encoding sulfur dioxygenase (*Sdo*) was present, which oxidizes S^0^ to sulfite in *A. thiooxidans* (13). Several copies of the heterodisulfide reductase complex genes (*hdrABC*) were present in the genome of strain RS19-104. In lithotrophic sulfur oxidizers, HdrABC is thought to oxidize S^0^ to sulfite with the assistance of rhodanese, DsrE, and TusA-like sulfurtransferases (14). The Hdr complex operon found in *Acidithiobacillus ferrooxidans* is also conserved in several acidophilic sulfur-oxidizing taxa (15, 16) as well as in strain RS19-104. Unlike HdrA in methanogenic archaea, HdrA in sulfur oxidizers is thought to be non-electron-bifurcating because it lacks central and carboxy-terminal ferredoxin-binding domains as well as an N terminal [4Fe-4S] cluster-binding domain (14). These domains were also absent from HdrA in strain RS19-104. If it is unable to bind and accept electrons from ferredoxin, HdrA may accept electrons from the quinone pool to reduce sulfite to S^0^.

It is also possible that sulfate could be reduced directly to S^0^ under comproportionation conditions. Supported by isotopic labeling, anaerobic methanotrophic (ANME) archaea have been shown to perform direct reduction of sulfate to S^0^ under extremely reducing conditions (17). Because sulfate reduction genes were absent in the genome, it was proposed that sulfate reduction by ANME archaea was catalyzed by a novel enzymatic pathway. Although this novel pathway has not yet been verified, a similar pathway could also be used by comproportionating microbes to produce S^0^ from sulfate.

In our proposed comproportionation pathway, sulfide is oxidized via Sqr to S^0^, donating two electrons to the quinone pool (Fig. 2). Sulfate is reduced either to sulfite by the assimilatory sulfate reduction pathway or to S^0^ by a novel metabolic pathway. The sulfite produced by sulfate reduction is reduced to S^0^ by reverse-acting Sdo, HdrABC, or SoeABC.

**Figure 2.**
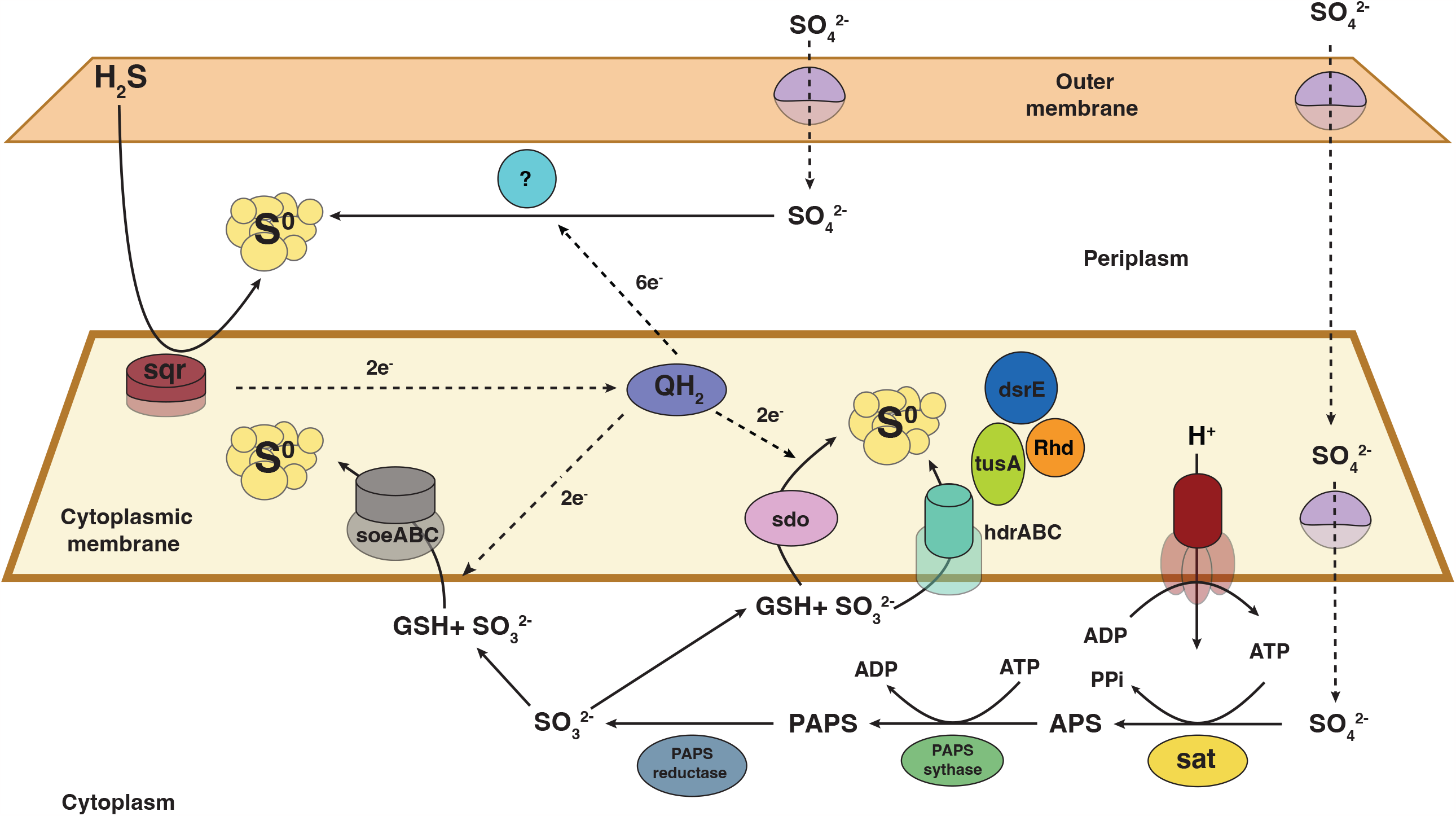
Proposed comproportionation pathways of strain RS19-104 based on genes present in the whole genome sequence. The blue circle with a question mark indicates an unknown enzyme that could catalyze the direct reduction of sulfate to S^0^. Gene names can be found in Table 3. GSH represents glutathione.

Although strain RS19-104 grew under comproportionation conditions for several transfers and we obtained enough biomass for whole genome sequencing, we were unable to maintain the culture growth for long periods. The culture was ultimately lost, but the experimental framework outlined here can be applied to future searches for sulfur comproportionators in other natural and impacted environments with favorable conditions, including extreme acid mine drainage pit lakes, certain terrestrial volcanic springs and shallow marine vents, subglacial lakes, and acid-sulfate crater lakes. In addition to measuring consumption of sulfate and sulfide and the production of S^0^, multi-isotope incubations should be conducted to show that S atoms from sulfate and sulfide are incorporated into S^0^ produced during cell growth (Supplementary methods). Under aerobic sulfide oxidation conditions, *A. thiooxidans* accumulates S^0^ inclusions that can be observed by TEM (18). If comproportionation produces similar intracellular S^0^, isotopic label from sulfate and sulfide could be detected in intracellular inclusions via NanoSIMS. Additionally, because of the high background concentration of sulfate and sulfide in the comproportionation culture medium, radioisotope incubations may be advantageous for measuring reactant consumption rates. Future searches for sulfur comproportionators could use a genomics-guided approach to cultivation. Metagenome-assembled genomes from target environments that have both sulfide oxidation and dissimilatory sulfate reduction pathways could be targeted for cultivation.

## Materials and Methods

### Sample collection and cultivation

Approximately 0.5 cm^3^ of snottite was collected from Ramo Sulfureo in the Frasassi cave system in July 2019. Snottite pH was measured in the field with pH paper (range 0 – 2.5). H_2_S_(g)_ concentrations were measured above the stream using an ENMET RECON/4a portable gas detector. Snottites were removed from the cave wall using sterile forceps and placed into microcentrifuge tubes containing comproportionation cultivation medium. A sterile syringe was used to transfer the snottite-medium slurry into a serum bottle containing 50 mL medium and sealed with a blue butyl stopper. Inoculated bottles were incubated at 15ºC until return to USC. The culture was incubated at 15ºC and purified by dilution to extinction. Purity of the resulting isolate was confirmed by Sanger sequencing of the 16S rRNA gene.

### Cell Counting

Samples were fixed with glutaraldehyde (2.5% final concentration), incubated at room temperature for 1 hour, neutralized with 5M NaOH, and stained with 10X SYBR Green I. Samples were filtered through black polycarbonate filters and visualized under a fluorescence microscope.

### DNA extraction and bioinformatics

Samples for DNA extraction were collected by filtering 2 mL enrichment medium through a sterile 25 mm, 0.1 μm Supor filter. DNA for 16S rRNA gene sequencing was extracted from filters using the Qiagen PowerBiofilm DNA extraction kit following manufacturer instructions. Full-length 16S rRNA genes were amplified from isolate cultures using the 27F/1492R primer pair. Sanger sequencing was performed by GeneWiz (La Jolla, CA). DNA for whole genome sequencing was extracted from filters using a combination of proteinase K digestion, phenol-chloroform-isoamyl alcohol extraction, and ethanol precipitation (https://www.protocols.io/view/modified-phenol-chloroform-genomic-dna-extraction-e6nvwkjzwvmk/v2). Whole genome sequencing was performed at the Caltech Genetics and Genomics Laboratory. Oxford Nanopore sequencing libraries were constructed using the PCR Barcoding Kit (SQK-PBK004) and were sequenced using MinION flowcells (FLO-MIN106). Basecalling was performed with ONT Guppy v3.4.5. Illumina libraries were constructed using the NEBNext Ultra kit. Single end, 100 basepair reads were sequenced using HiSeq2500. The 16S rRNA sequence and whole genome sequence are available in GenBank under BioProject accession number XXXXXX.

### Thermodynamic calculations

The Gibbs energy yields (Δ*G*_*r*_) for sulfur comproportionation and sulfate reduction coupled to ammonium oxidation were calculated as described in Alain *et al*. 2022 (22).

## Acknowledgements

We thank A. Montanari for providing logistical support and use of facilities and laboratory space at the Osservatorio Geologico di Coldigioco, Apiro, Italy. Thanks to the members of the Gruppo Speleologico

C.A.I. di Fabriano for technical assistance during field campaigns, and to Dan Jones, Zoë Havlena, and Mackenzie Best for assistance in the field. We also thank Fabai Wu and Igor Antoshechkin for assistance with whole genome sequencing. HSA was supported by an NSF Graduate Research Fellowship grant DGE-1842487, the Lewis and Clark Fund for Exploration and Field Research in Astrobiology, the National Cave and Karst Institute NCKRI Scholar Fellowship, the Josephine de Karman Fellowship, and the Cave Conservancy Foundation Graduate Fellowship in Karst Studies.

## Supplementary methods

### Cultivation

Custom cultivation medium for sulfur comproportionation was prepared under 80% N_2_-20% CO_2_. The basal medium contained the following (per liter of 50 mM sulfuric acid): 2.42 g Na_2_SO_4_, 0.50 g MgSO_4_·7H_2_O, 0.025 g CaCl_2_·2H_2_O, 0.1 g KCl, 0.13 g (NH_4_)_2_SO_4_, 0.14 g Na_2_HPO_4_, 1 mL trace element solution, and 0.5 mL of 0.2% resazurin. The pH was adjusted to 1 with concentrated sulfuric acid and the medium was dispensed into serum bottles. Bottles were stoppered and crimped, and a slight overpressure was added to the headspace with 80% N_2_-20% CO_2_. After autoclaving, the medium was amended with 1 mL of vitamin solution, sodium bicarbonate at a final concentration of 10 mM, and Na_2_S·9H_2_O at a final concentration of 2 mM. The trace element contained the following (per liter of water): 1.7 mL 20 mM HCl, 2.10 g FeSO_4_·7H_2_O, 0.03 g H_3_BO_3_, 0.10 g MnCl_2_·4H_2_O, 0.19 g CoCl_2_·6H_2_O, 0.024 g NiCl_2_·6H_2_O, 0.002 g CuCl_2_·6H_2_O, 0.144 g ZnSO_4_·7H_2_O, 0.036 g Na_2_MoO_4_·2H_2_O, 0.0326 g VOSO_4_·H_2_O, 0.025 Na_2_WO_4_·2H_2_O, 0.006 g Na_2_SeO_3_·5H_2_O. The vitamin solution contained the following (per liter 10 mM MOPS, pH 7.2): 0.10 g riboflavin, 0.03 g biotin, 0.10 g thiamine HCl, 0.10 g L-ascorbic acid, 0.10 g D-calcium pantothenate, 0.10 g folic acid, 0.10 g nicotinic acid, 0.10 g 4-aminobenzoic acid, 0.10 g pyridoxine HCl, 0.10 g lipoic acid, 0.10 g thiamine pyrophosphate, 0.01 g cyanocobalamin.

### Bioinformatics

Nanopore long reads were assembled using Canu v.2.1.1 (1). Reads were polished with Racon v1.4.20 (2) and Medaka v1.1.1 (https://github.com/nanoporetech/medaka). Reads with Q scores below 7 were filtered and barcodes and adapters were trimmed. The resulting assembly was mapped using BamM (http://ecogenomics.github.io/BamM/) with 150 bp Illumina paired-end reads. Mapped reads were used for error correction using Pilon v1.2.2 (3). Genome quality was determined using CheckM2 v1.0.1 (4). Closely related strains were identified using GToTree (5). The command “gtt-get-accessions-from-GTDB” was used to collect representative genomes from the genus *Acidithiobacillus* that were present in the Genome Taxonomy Database (6), and the command “GToTree” was used to construct a de novo phylogenomic tree of 172 Gammaproteobacterial single copy genes. Average nucleotide identity (ANI) and average amino acid identity (AAI) were calculated for all genomes in the *Acidithiobacillus* genus using the IMG ANIcalculator (7) and EzAAI (8). Metabolic functional traits were predicted using METABOLIC v4.0 (9), MetaSanity v1.3.0 (10), eggNOG-mapper v2 (11), and SCycDB (12).

### Thermodynamic calculations

The Gibbs energy yields (Δ*G*_*r*_) for sulfur comproportionation and sulfate reduction coupled to ammonium oxidation were calculated with

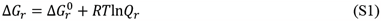

where Δ*G*_*r*_^0^ is the standard state Gibbs energy, *R* is the universal gas constant, T is the temperature in Kelvin, and *Q*_*r*_ refers to the reaction activity quotient. Values of Δ*G*_*r*_^0^ were calculated at *in situ* or media temperatures and 1 bar with the revised Helgeson-Kirkham-Flowers (HKF) equations of state (13–15) using the “subcrt” command from the R software package CHNOSZ v1.4.1 (16). Thermodynamic data in CHNOSZ are derived from the OrganoBioGeoTherm database, which come from a number of sources (https://chnosz.net/download/refs.html). The sources of these data are provided in the CHNOSZ package documentation. Values of *Q*_*r*_ were calculated with

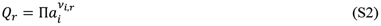

where *a*_*i*_ represents the activity of the *i*^th^ species raised to its stochiometric reaction coefficient ν_*i,r*_, in the *r*^th^ reaction, which is positive for products and negative for reactants. Activities were calculated with the relation

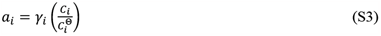

where *γ*_*i*_ and *C*_*i*_ are the activity coefficient and concentration of the *i*^th^ species, respectively. *C*_*i*_^Θ^ is the concentration of the *i*^th^ species under standard state conditions, which is equal to one molal referenced to infinite dilution. Activities for aqueous species were determined using the aqueous speciation package AqEquil v0.9.1 (17), which is based on EQ3/6 (18). The activities of S^0^ and water were taken to be unity (*a*_*i*_ = 1). Concentrations were sourced from the sulfur comproportionation medium recipe and from *in situ* geochemical concentrations. H_2_S_(g)_ concentrations in the cave air ranged from 15 to 33 ppmv. Sulfate concentration in the snottite was inferred from the pH (0 -1).

The Gibbs energy for sulfate reduction coupled to ammonium oxidation was calculated under medium conditions at 15ºC for six reactions with different S and N products:

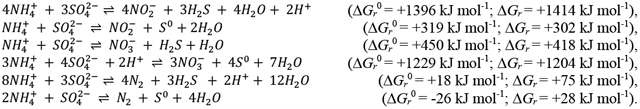

### Chemical measurements

Sulfate, sulfide, and S^0^ should be measured in comproportionation cultures using ion chromatography, the Cline assay (19), and high performance liquid chromatography, respectively. A method for extracting and measuring S^0^ is described in McGuire and Hamers 2000 (20)

### Multi-isotope probing and nanoSIMS

Multi-isotope incubations were performed with strain RS19-104 using 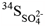 (90 atom% ^34^S, Sigma Aldrich) and 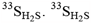 was prepared in house by the reduction of ^33^S^0^ (99%, Sigma Aldrich). Fe^33^S was synthesized from ^33^S^0^ and carbonyl iron (Sigma Aldrich, ≥97% Fe basis). ^33^S^0^ and an excess of carbonyl iron were mixed in a Hungate tube sealed with a butyl rubber stopper and the headspace was flushed for 30 minutes with N_2_. The mixture was then ignited with an ethanol burner flame. The Fe^33^S was placed into a 150 mL serum bottle sealed with a butyl rubber stopper and the headspace was flushed for 30 minutes with N_2_, then placed on ice. 6.0 M HCl was sparged for 20 minutes with N_2_ on ice. To form 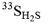 HCl was added with an N_2_-sparged syringe to the serum bottle containing Fe^33^S. The bottle was removed from the ice bath after 10 minutes and incubated at room temperature overnight. The following day, the bottle was placed in a sonicator bath to dissolve remaining Fe^33^S particles. Cold distillation of 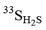 was performed by flushing the bottle containing 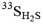 with N_2_ for 10-15 minutes through 1.0 mM NaOH to trap NaH^33^S. The NaH^33^S was neutralized with HCl to form 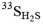. Conversion to H_2_S was verified and quantified with the Cline assay (19).

Modified comproportionation medium was prepared with 30 mM H_2_SO_4_ (instead of 50 mM) without the addition of Na_2_SO_4_ and with only 0.5 mM (NH_4_)_2_SO_4_. 10 mL of medium was dispensed into Balch tubes under 80% N_2_-20% CO_2_, and the tubes were stoppered and autoclaved. After autoclaving, the medium was amended with sodium bicarbonate at a final concentration of 5 mM and Na_2_S x 9H_2_O at a final concentration of 1.6 mM. The following isotopically labeled compounds were added: 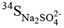-(final concentration 20 mM; 40 at. %), 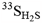 (final concentration 0.4 mM; 20 at. %), 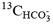 (final concentration 5 mM; 50 at. %), and 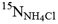 (final concentration 0.5 mM; 50 at. %). The at. % added to incubations was calculated by isotope mass balance: ^n^F_final_ = [(^n^F_unlabeled_ x m_unlabeled_) + (^n^F_labeled_ x m_labeled_)]/m_final_, where mass (m) was the amount of Na_2_SO_4_, H_2_S, NH_4_Cl, or HCO_3-_ added to the mixture and at. % = 100 x ^n^F. The final medium was inoculated sulfur comproportionation cultures. After 1 month of incubation at 15ºC, samples were fixed with glutaraldehyde (final concentration 2.5%), filtered onto a 0.1 μm, 13 mm Supor filter, dehydrated in an ethanol series, and coated with 20 nm gold. Carbon, nitrogen, and sulfur isotopic compositions were measured using a NanoSIMS 50L (CAMECA, Gennevilliers, France) housed at the Center for Microanalysis in the Division of Geological and Planetary Sciences at the California Institute of Technology. Cells were analyzed using a ∼4 pA primary Cs^+^ beam current. Seven masses were collected in parallel (^12^C, ^13^C, ^14^N, ^15^N, ^32^S, ^33^S, and ^34^S).

### Radioisotope and stable isotope probing

Strain RS19-104 was used for radioisotope incubations using 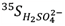 (Perkin Elmer) and 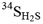 (synthesized as described above from ^34^S – Sigma Aldrich). 50 mL bottles of complete comproportionation medium were spiked with 1μCi 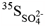 and 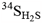 (final concentration 1 mM, 50 at. %). After 24 hours of incubation at 15 mL, a 25 mL sample was taken for sulfur pool separation. 5 mL of medium was filtered through a 0.1 μm, 25 mm Supor filter to capture cells. The filters were then added to a scintillation vial and covered with 4.5 mL Ultima Gold scintillation cocktail (Perkin Elmer). Sulfate, sulfide, and S^0^ pools were separated. 50 mL of 5M NaOH was added to a 2 mL microcentrifuge tube in triplicate. With a syringe, 0.8 mL of the culture medium and 0.8 mL of 20% ZnCl_2_ was added to each of the 2 mL tubes to precipitate ZnS. The tubes were vortexed and centrifuged at 10,000 rpm for 10 minutes. The supernatant was removed and transferred to new tubes. 400 μL of 1M BaCl_2_ was added to the supernatant to precipitate BaSO_4_. The tubes were vortex and centrifuged at 10,000 rpm for 10 minutes. The supernatant was removed and transferred to a scintillation vial. 2 mL of tetrachloroethylene was added to extract S^0^ (20) and the scintillation vials were shaken overnight. The following day, the organic layer was removed from the scintillation vial and transferred to a new vial. Both the aqueous and organic layers were covered with 4.5 mL scintillation cocktail. The precipitated ZnS and BaSO_4_ were resuspended in 1 mL diH_2_O and split into two scintillation vials. 4.5 mL Ultima Gold was added to one vial and the other was frozen for ^34^S measurements. Scintillation counting was performed on a Beckman Coulter LS 6000 Scintillation System.

